# Targeting MLL methyltransferases enhance the anti-tumor effects of PI3K inhibition in hormone receptor-positive breast cancer

**DOI:** 10.1101/2021.10.01.462826

**Authors:** RB. Jones, J. Farhi, M. Adams, K. Parwani, M. Zecevic, G. Cooper, RS. Lee, AL. Hong, JM Spangle

## Abstract

The high frequency of aberrant PI3K pathway activation in hormone receptor-positive (HR+) breast cancer has led to the development, clinical testing, and approval of the p110α-selective PI3K inhibitor alpelisib. The limited clinical efficacy of alpelisib and other PI3K inhibitors is partially attributed to the functional antagonism between PI3K and estrogen receptor (ER) signaling, which is mitigated via combined PI3K inhibition and endocrine therapy. We and others have previously demonstrated a chromatin-associated mechanism by which PI3K supports cancer development and antagonizes ER signaling through the modulation of H3K4 methylation. Here we show that inhibition of the H3K4 histone methyltransferase MLL1 in combination with PI3K inhibition impairs HR+ breast cancer clonogenicity and cell proliferation. While combined PI3K/MLL1 inhibition reduces AKT effector signaling and H3K4 methylation, MLL1 inhibition increases PI3K effector signaling and upregulates the expression of receptor tyrosine kinase signaling cascades upstream of and including AKT. These data reveal a feedback loop between MLL1 and AKT in which MLL1 inhibition reactivates AKT. We additionally show that combined PI3K and MLL1 inhibition synergizes to cause cell death in *in vitro* and *in vivo* models of HR+ breast cancer, which is enhanced by the additional genetic ablation of the H3K4 methyltransferase and AKT target MLL4. Together, our data provide evidence of a feedback mechanism connecting histone methylation with AKT and may support the preclinical development and testing of pan-MLL inhibitors.

**Significance Statement:** Pharmacological inhibition of PI3K provides limited efficacy in PIK3CA-mutated, HR+ breast cancers. Here the authors leverage PI3K/AKT-driven chromatin modification to identify MLL histone methyltransferases as a therapeutic target. Dual PI3K and MLL inhibition synergize to reduce clonogenicity and cell proliferation while enhancing apoptosis in *in vitro* models, and induces tumor regression in xenograft models of PI3K-activated, HR+ breast cancer. Furthermore, MLL1 inhibition reveals a feedback loop leading to AKT hyperactivation, which is relieved with combined PI3K/MLL inhibition. These findings demonstrate the utility of MLL inhibitors for the treatment of some solid cancers, as patients with HR+ breast cancer characterized by PIK3CA mutation may derive clinical benefit from combined PI3K/MLL inhibition.

## Introduction

The Phosphatidylinositol-3-kinase (PI3K) signaling pathway coordinates cellular response to external stimuli, supporting cell growth, metabolism, proliferation, and survival, cellular processes that are essential to cancer development and progression that can dictate therapeutic response^1^. Class IA PI3Ks are heterodimeric protein complexes that consist of a catalytic p110 isoform and a regulatory p85 isoform^2^. The genes PIK3CA, PIK3CB, and PIK3CD encode the catalytic subunit of the p110α, p110β, and p110δ PI3Ks, respectively, which complex with regulatory p85 isoforms^1^. Activated PI3Ks catalyze PIP2 into PIP3, which leads to activation of AKT and other downstream effectors. While PI3K activity supports biological processes mediated independently of AKT, AKT is considered a critical and principal PI3K effector^3^. With more than 200 identified substrates, AKT is a critical signaling amplifier that integrates upstream PI3K signaling with downstream biological functions through substrate phosphorylation^3^. Aberrant PI3K/AKT hyperactivation is common in human cancers and occurs via multiple distinct genomic alterations, e.g., receptor tyrosine kinase (RTK) mutation or amplification, PTEN functional inactivation via deletion or mutation, PIK3R1 or AKT1 mutation. Collectively, PIK3CA mutational activation is among the most common oncogenic driver events in breast cancer^1,4^. PIK3CA mutation renders the p110α protein product constitutively active, leading to AKT hyperactivation^5^.

The importance of cancer-associated genomic alterations, including PIK3CA mutation in the PI3K pathway and its subsequent dysregulation, has prompted the development and clinical testing of numerous PI3K and AKT inhibitors^6^. Given the divergent functions of PI3K isoforms, isoform-selective PI3K inhibitors exhibit enhanced preclinical and clinical success compared to pan-PI3K agents due to the on-target selectivity and has resulted in the FDA approval of the several isoform-selective PI3K inhibitors: idelalisib^7^, copanlisib^8^, and alpelisib^9^. Alpelisib is a p110α-selective inhibitor, approved for the treatment of metastatic PIK3CA-mutant, HR+ breast cancers in combination with the estrogen receptor antagonist fulvestrant^9^. Despite this success, preclinical and clinical studies utilizing PI3K inhibitors including alpelisib demonstrate a lack of efficacy when used as monotherapies, and resistance remains a common occurrence. Thus, achieving an efficacious response to PI3K inhibition remains an unmet clinical need.

Histones, including H3, serve an integral role downstream of signaling pathways by supporting a variety of posttranslational modifications (PTMs) including phosphorylation, acetylation, and methylation^10^. The precise combination of histone PTMs contribute to chromatin state and ultimately gene expression. Evidence suggests that some enzymes responsible for the reading, writing, and erasure of histone PTMs are effectors of signaling cascades including PI3K. AKT has been shown to phosphorylate the H3K27 histone methyltransferase EZH2^11^ and the histone acetyltransferase p300^12^. Breast cancer-associated redundant phosphorylation of the H3K4 demethylase KDM5A^13^ and the H3K4 methyltransferase MLL4/KMT2D^14^ by AKT modulates H3K4 methylation and enhances gene expression to support pro-oncogenic growth and therapeutic resistance, indicating that H3K4 methylation and associated epigenomic/cistromic regulation may be a critical downstream effector of aberrant PI3K/AKT signaling in breast and other cancers. Clinical studies including metaanalyses demonstrate that aggressive breast and other cancers are characterized by elevated H3K4 methylation and that H3K4 trimethylation (H3K4me3) is an indicator of poor patient prognosis^15–17^.

Here, we demonstrate that combined therapeutic targeting of PI3K/AKT and the H3K4 methyltransferase MLL1 reduces PI3K effector phosphorylation and H3K4 methylation in PIK3CA-mutated HR+ breast cancer cell lines. Dual PI3K and MLL1 inhibition reduces breast cancer cell clonogenicity and synergizes to reduce cell viability and increase apoptosis, effects that are further enhanced following genetic loss of MLL4. We also uncover a previously uncharacterized feedback loop between MLL and AKT, demonstrating that pharmacological MLL inhibition relieves PI3K repression and elevates AKT activity, and combined PI3K and MLL1 inhibition abrogate aberrant AKT and effector activation.

## Results

### Combined PI3K/MLL1 inhibition reduces breast cancer cell line clonogenicity through on-target activity

Previous reports demonstrate that aberrant PI3K/AKT signaling regulates H3K4 methylation through the activity of H3K4-enzymes in hormone receptor-positive (HR+) breast cancer^13,14^ and high H3K4 methylation is associated with poor prognosis in breast and other cancers^15^. We thus postulated that combined PI3K and MLL inhibition may provide additional therapeutic benefit in PI3K pathway activated, HR+ breast cancer. To assess the effects of combined PI3K/MLL1 inhibition on cell growth, the PI3K-activated (PIK3CA^H1047R^-mutant), HR+ breast cancer cell line MCF7 was treated with the PI3K inhibitor alpelisib, the MLL1 inhibitor MI-503, or the combination. Combined PI3K and MLL1 inhibition reduced the clonogenicity and cell proliferation compared to either monotherapy (Fig 1A), which was observed at multiple doses (Fig S1A). MI-503 is a small-molecule MLL1 inhibitor that disrupts the association between MLL1 and its co-factor Menin, thereby disrupting target gene expresion^18^. Cells treated with the MLL-Menin inhibitor MI-136^19^ in combination with PI3K inhibitors also exhibit impaired clonogenicity (Fig 1A). A reduction in clonogenicity was also observed in cells treated with MLL1 inhibitors in combination with the p110β-sparing PI3K inhibitor taselisib^20^ (Fig 1B) or the pan-PI3K inhibitor pictilisib^21^ (Fig S1B). Combined PI3K and MLL1 inhibition also decreased clonogenicity in the HR+, PIK3CA^H1047R^-mutant cell line T47D (Fig 1C, D, Fig S1C). These data suggest that combined PI3K and MLL1 inhibition impairs the clonogenicity in PI3K-activated, HR+, breast cancer models.

**Figure 1.**
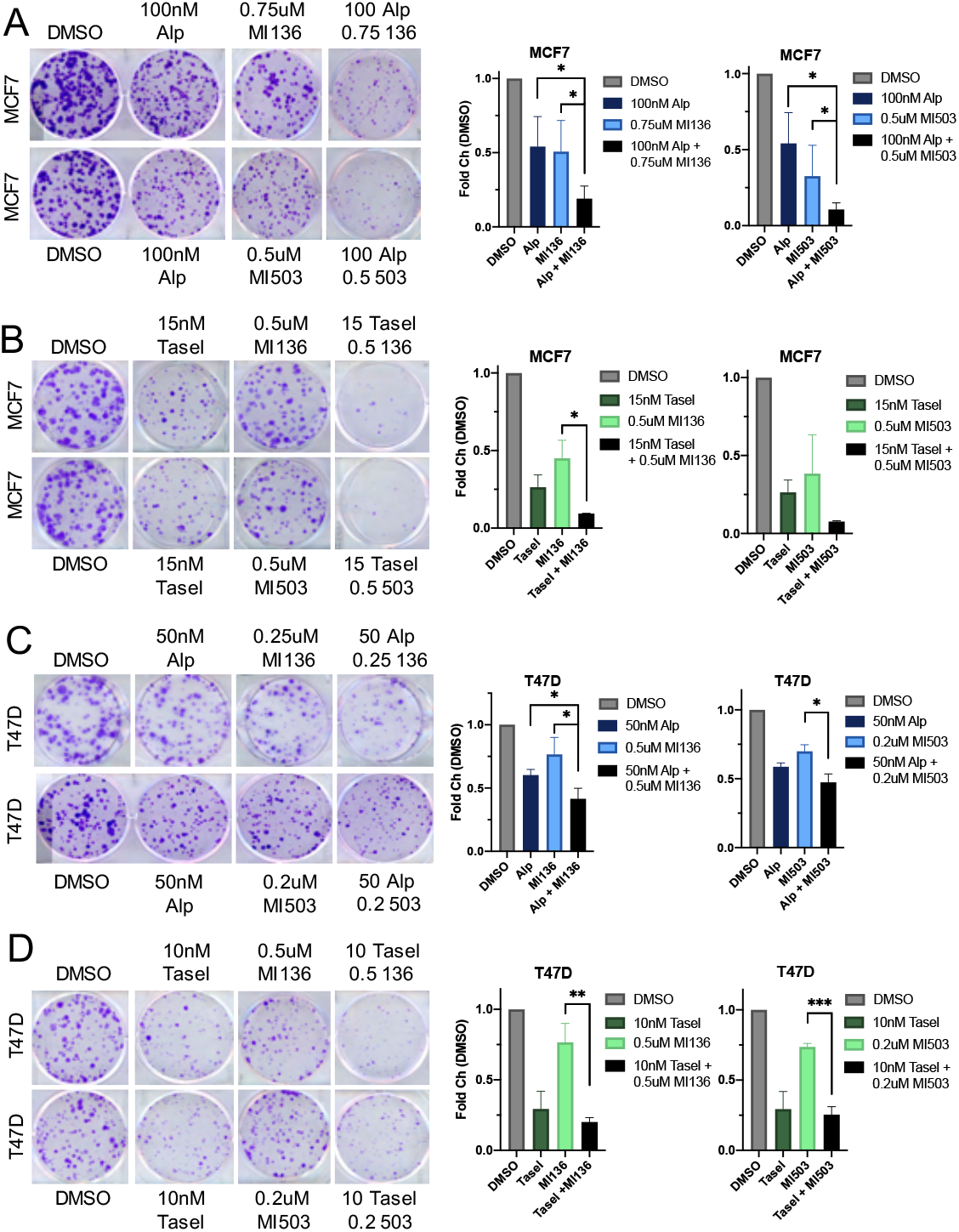
Combined PI3K and MLL1 inhibition reduces clonogenicity of HR+, PIK3CA-mutant, breast cancers. (A) MCF7 breast cancer cells treated with the indicated concentrations of alpelisib, MI-136, MI-503, or DMSO for 19 d before fixation and crystal violet staining. (B) MCF7 breast cancer cells treated with the indicated concentrations of taselisib, MI136, MI-503, or DMSO for 19 d before fixation and crystal violet staining. (C) T47D breast cancer cells treated with the indicated concentrations of alpelisib, MI-136, MI-503, or DMSO for 19 d before fixation and crystal violet staining. (D) T47D breast cancer cells treated with the indicated concentrations of taselisib, MI-136, MI-503, or DMSO for 19 d before fixation and crystal violet staining. For all experiments, representative images from at least 3 independent experiments are shown. Data are shown as mean ± SEM. \**P* < 0.05; \*\**P* < 0.01; \*\*\**P* < 0.001.

We next determined whether the biological activity of PI3K and MLL1 inhibitors observed (Fig 1) is a result of on-target enzymatic inhibition. MCF7 and T47D cells were treated with PI3K and/or MLL1 inhibitors, and PI3K inhibition with the p110α-selective inhibitor alpelisib reduced effector signaling as indicated by a decrease in phosphorylated AKT and S6 (Fig 2A) and MLL inhibition via MI-503 or MI-136 modestly reduced H3K4 methylation (Fig 2A). Similar results were observed with the p110β-sparing PI3K inhibitor taselisib and the pan-PI3K inhibitor pictilisib (Fig S2A). Combined PI3K and MLL1 inhibition reduced H3K4 methylation and was detectable as early as 72h (Fig 2A). Robust MLL1-inhibitor monotherapy – directed loss of H3K4 methylation was detected at later time points such as 96-120h (Fig S2B), which is consistent for detecting global changes to histone PTMs in response to therapeutic inhibition^13^.

**Figure 2:**
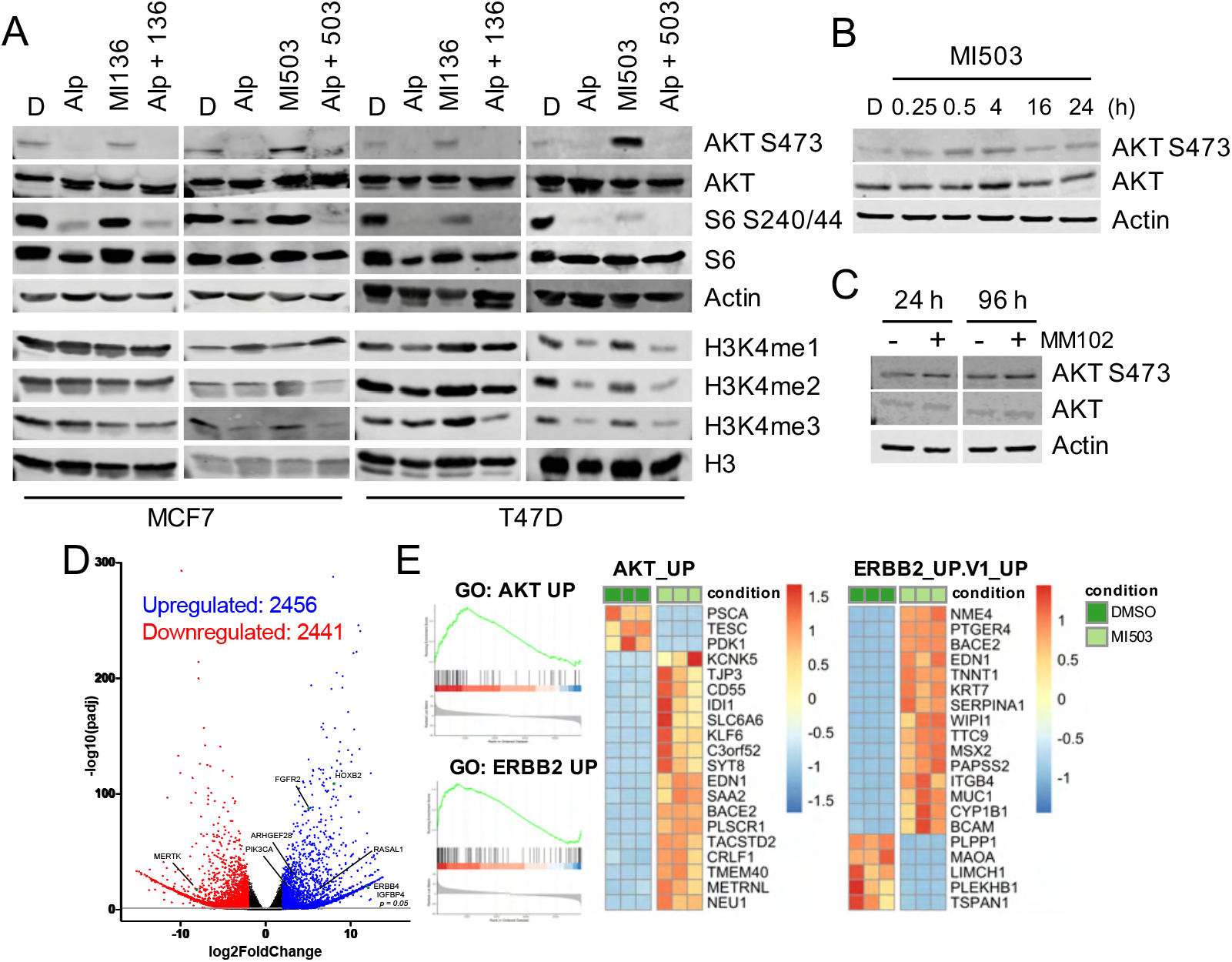
MLL1 inhibition hyperactivates AKT through a feedback loop. (A) MCF7 or T47D breast cancer cells were treated with alpelisib (1 uM), taselisib (0.5 uM), MI-136 (4 uM), MI-503 (4 uM) or DMSO for 24h (top, PI3K effector) or 72h (bottom, histone modifications) prior to protein extraction. Lysates were immunoblotted for the indicated antibodies. (B) T47D cells treated with the MI-503 (4 uM) for the indicated time and then lysates prepared. Lysates were immunoblotted for the indicated antibodies. (C) T47D cells treated with MM102 (6 uM) for the indicated time and then lysates prepared. Lysates were immunoblotted for the indicated antibodies. (D) Volcano plot of statistical significance *(P* = 0.05) against fold change between T47D cells treated with control (DMSO) and MI-503-(4 uM) for 24h. (E) GSEA depicting GO classifications of differentially expressed gene sets between the DMSO and MI-503 cohorts shown in (D). (F) Heat maps representing the top 20 differentially expressed genes from GO classifications shown in (E).

### MLL inhibition induces feedback activation of AKT signaling

Unexpectedly, we found that MLL inhibition in PI3K-activated cancer cell line models increases PI3K effector signaling as indicated by increased AKT and S6 phosphorylation, which was abrogated with combined PI3K and MLL inhibition (Fig 2A, Fig S2A, B). The detected increase in PI3K effector signaling following MLL1 monotherapy suggests the presence of a feedback loop between MLL and AKT. Similar MLL1 inhibitor-induced PI3K effector activity was detected in the PTEN-null prostate cancer cell line LNCAP, which is also characterized by PI3K pathway hyperactivation (data not shown). We found that MLL 1 inhibition increased AKT phosphorylation as early as 5min post inhibition, which was sustained for at least 24h (Fig 2B). To test whether the ability of MLL1 inhibition to enhance PI3K effector activity is due to on-target inhibitor activity, we treated T47D cells with a panel of MLL1 inhibitors that function through disparate mechanisms of action and found that MLL1/Menin (MI-503 and MI-136) and MLL/WDR5 (MM102) inhibitors increased AKT phosphorylation (Fig 2C). Collectively, these data suggest a functional relationship between PI3K/AKT signaling and MLL methyltransferases which may support their combined inhibition. To determine whether MLL inhibition may remodel chromatin to subsequently alter gene expression to support AKT activation, we performed RNA-seq on T47D cells treated with the MLL inhibitor MI-503 for 24h. Expression of more than 2400 genes was upregulated following MLL inhibition, including genes involved in upstream receptor tyrosine kinase signaling (FGFR2, ERBB4, IGFBP4), as well as PI3K/Ras signaling (PIK3CA) (Fig 2D). MLL inhibition significantly dysregulated the expression of biological pathways and processes, including upregulation of ERBB2, RAS, RAF, mTOR, and AKT signaling, which promote AKT and AKT effector activation (Fig S3A). Gene set enrichment analysis (GSEA) further demonstrated that treatment with MLL inhibitors significantly upregulates the expression of AKT and ERBB2 pathway genes (Fig 2E, S3B). Collectively, these data demonstrate MLL inhibition promotes the activation of AKT signaling through the presence of feedback loop between MLL and upstream of AKT.

### Combined PI3K/MLL1 inhibition synergizes to reduce viability by enhancing apoptosis in vitro

A major limitation of clinical PI3K inhibition is the lack of apoptotic induction that leads to tumor clearance, as PI3K monotherapy is primarily associated with a cytostatic effect induced by G1 arrest^22^. To determine whether combined PI3K/MLL1 inhibition overcomes PI3K inhibitor induced cytostasis, we treated PI3K-activated, HR+ breast cancer cells with the PI3K inhibitor alpelisib, the MLL1 inhibitor MI-503, or the combination, and cellular viability was evaluated. Combined PI3K and MLL inhibition reduced cell viability in multiple models of breast cancer (Fig 3A, B), which was confirmed with additional PI3K and MLL1 inhibitors (Fig 3A, B, Fig S3A). Dual PI3K/MLL inhibition selectively reduced the viability of PIK3CA-mutated, HR+ breast cancer cell lines, as treatment with PI3K and MLL inhibitors did not reduce viability of the normal breast cell line MCF10A (Fig S4B). We next aimed to determine whether PI3K and MLL1 combination therapy is synergistic, as synergistic regimens potentially maximize therapeutic effects while minimizing off-target effects^23^. Drug synergy was calculated using both Loewe and Bliss, which measure synergy within the same pathway or among distinct pathways, respectively. Bliss and Loewe calculations/scores separately defined the combination of low-dose PI3K and MLL1 inhibition as synergistic; synergy was not achieved for the normal breast cell line MCF10A (Fig 3C)^24^. While PI3K or MLL1 monotherapies do not yield apoptotic induction, we found that the combination of PI3K and MLL1 inhibition increased apoptosis (Fig 3D), which is consistent with the metabolism-oriented cell viability assays (Fig 3A). To assess whether the detected loss of viability is due to on-target effects of MLL1 inhibition, we utilized the MLL1 inhibitor MM102. While MM102 is not as potent as MI-503 or MI-136, the peptidomimetic MM102 inhibits MLL1 function through the disruption of the MLL1-WDR5 protein-protein interaction, which is essential for MLL1 histone methyltransferase activity^25^. Combined PI3K inhibition with MM102 reduces cell viability and proliferation in models of PI3K-activated breast cancers (Fig S3A). These data demonstrate that combined PI3K and MLL inhibition synergize to reduce viability in models of PI3K activated, HR+, breast cancer.

**Figure 3.**
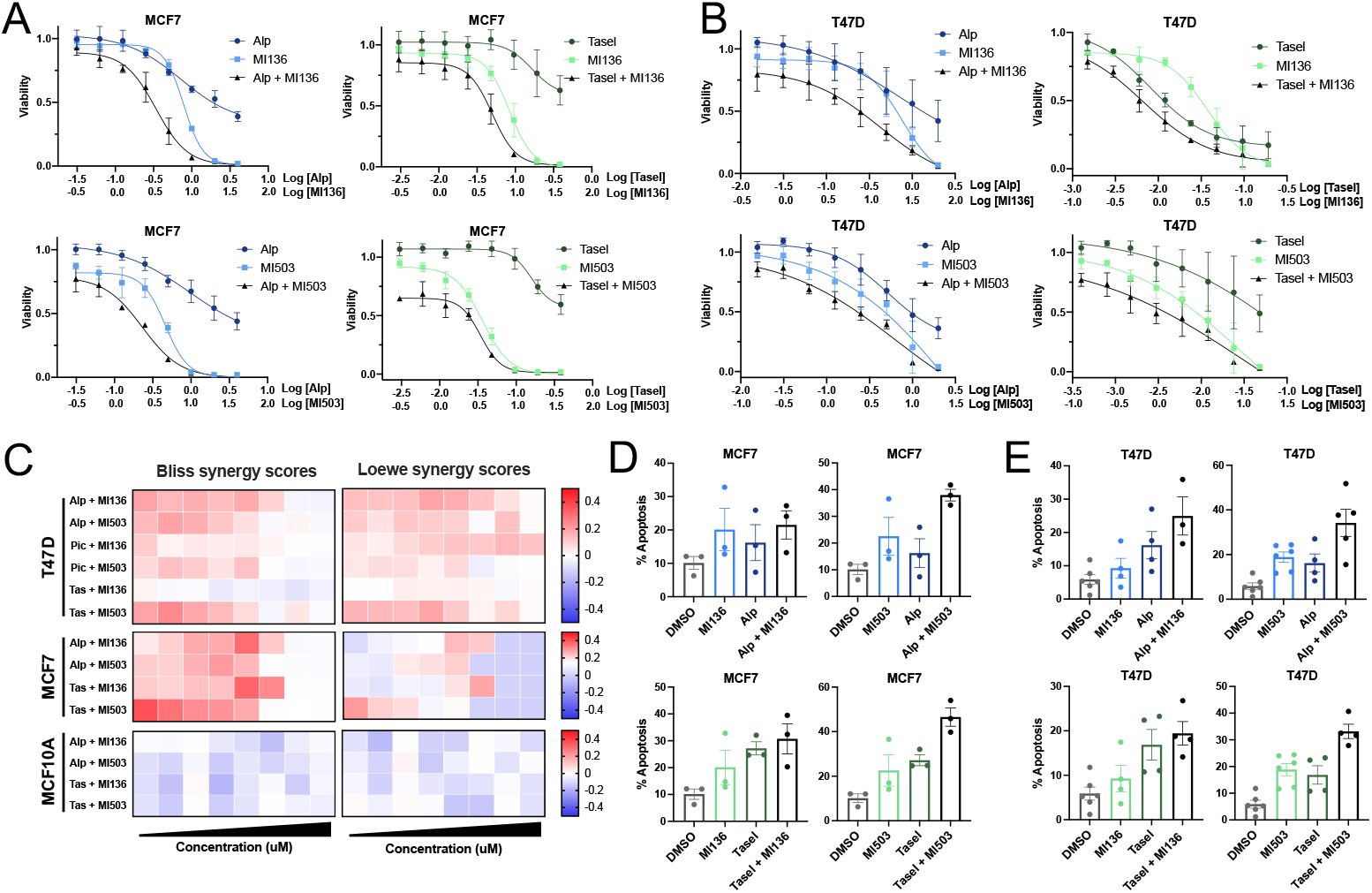
PI3K and MLL1 combination therapy synergizes to reduce viability and enhance apoptosis. (A) Cell viability curves in MCF7 breast cancer cells treated with an 8-point range of DMSO, alpelisib, taselisib, MI-503, and/or MI-136 for 120h. Results shown are representative of at least 3 independent experiments. Data are shown as mean ± SEM. (B) Cell viability curves in T47D breast cancer cells treated with an 8-point range of DMSO, alpelisib, taselisib, MI-503, and/or MI-136 for 120h. Results shown are representative of at least 3 independent experiments. Data are shown as mean ± SEM. (C) Bliss (left) and Loewe (right) synergy calculated from datapoints in (A), (B), and Sup Fig 4. (D) Annexin V staining in MCF7 cells treated with DMSO, alpelisib (1 uM), taselisib (0.5 uM), MI-136 (4 uM), MI-503 (4 uM) for 120h. Results shown are representative of at least 3 independent experiments. Data are shown as mean ± SEM. (E) Annexin V staining in MCF7 cells treated with DMSO, alpelisib (1 uM), taselisib (0.5 uM), MI-136 (4 uM), MI-503 (4 uM) for 120h. Results shown are representative of at least 3 independent experiments. Data are shown as mean ± SEM.

### Dual PI3K/MLL1 inhibition reduces tumor growth in xenograft models of PIK3CA-activated, HR+ breast cancer

To determine whether HR+ breast tumors characterized by PI3K pathway activation are sensitive to dual PI3K/MLL1 inhibition, MCF7 cells were transplanted into the mammary fat pads of nude mice. Palpable tumors were treated with vehicle, alpelisib, MI-503, or the combination. While alpelisib- and MI-503-treated tumors were smaller than vehicle-treated tumors, combined PI3K and MLL1 inhibition significantly reduced tumor volume (Fig 4A) and end-point tumor weight (Fig 4B), without toxicity-associated changes to animal weight during the treatment duration (Fig S4A). Dual PI3K and MLL1 inhibition reduced PI3K effector phosphorylation, H3K4 methylation, and cellular proliferation (Fig 4C, D). MLL1 inhibition via MI-503 monotherapy enhanced AKT phosphorylation in some tumors (Fig 4C, D), which was consistent with *in vitro* MLL1 inhibition (Fig 2, Fig S2). Taken together, our preclinical data suggest that HR+ breast xenografts characterized by PIK3CA mutation exhibit therapeutic benefit from combined PI3K and MLL1 inhibition that significantly exceeds PI3K or MI-503 monotherapy. Furthermore, patients with this subset of breast tumors may be suitable candidates for dual PI3K and MLL1 inhibition.

**Figure 4.**
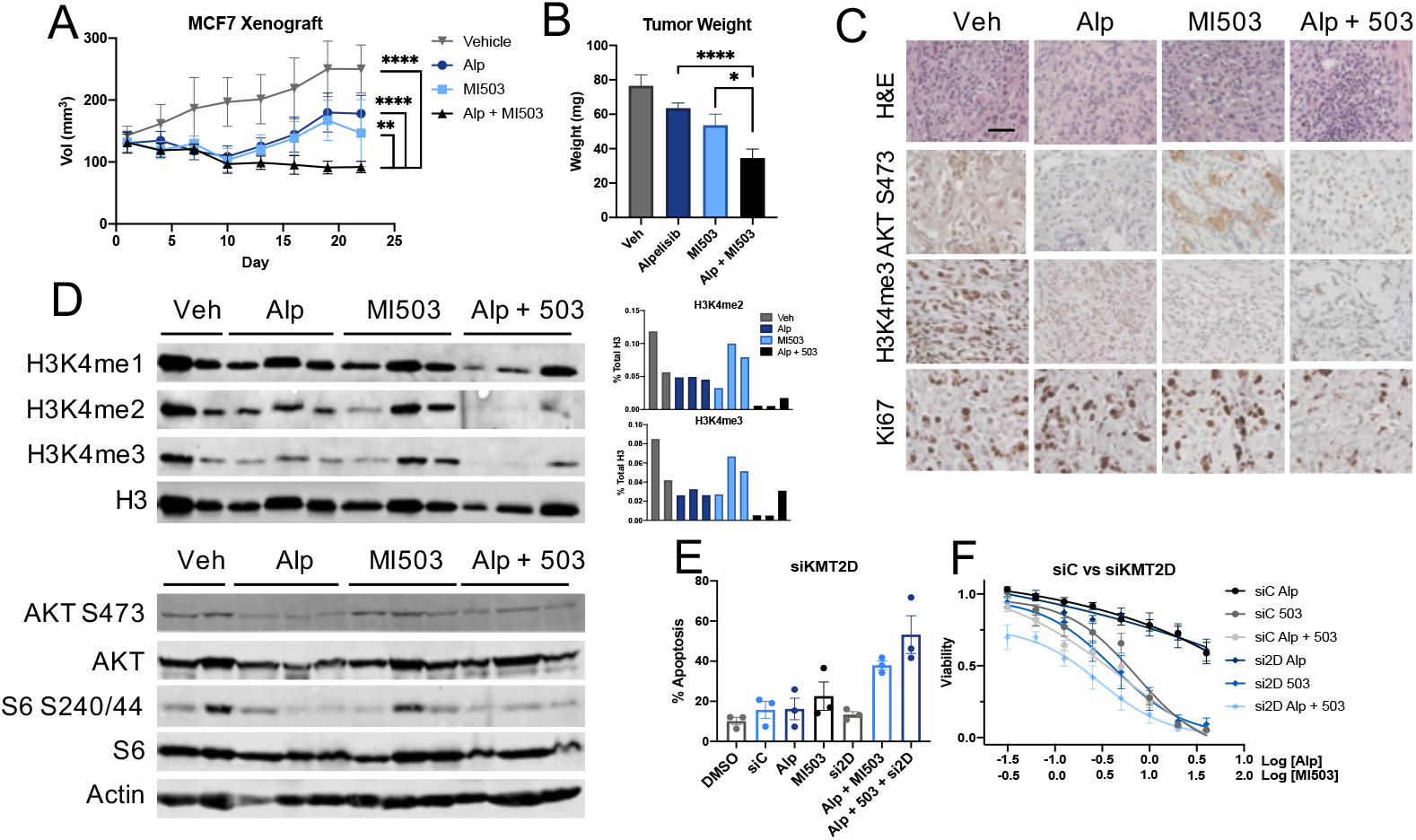
Combined MLL and PI3K inhibition provides therapeutic benefit in *in vivo* models of breast cancer. (A) Tumor volume of MCF7 cells implanted into the mammary fat pads of nude mice with once-daily treatment with alpelisib (35 mg/kg, gavage), MI-503 (30 mg/kg, IP), or the combination. End point tumor volume: MCF7 vehicle, 250.17 ± 11.24 mm^3^; MCF7 alpelisib, 178.11 ± 7.87 mm^3^; MCF7 MI-503, 146.59 ± 15.59 mm^3^; and MCF7 alpelisib/MI-503, 91.83 ± 2.69 mm^3^. Means ± SEM are shown; all groups *n* ≥ 12; two-way ANOVA with Tukey’s multiple-comparisons test. *\*\*P* < 0.01; \*\*\*\**P* < 0.0001. (B) The wet weight of individual mammary tumors plus associated mammary gland tissue from MCF7 vehicle (*n* = 12), MCF7 alpelisib (*n* = 13), MCF7 MI-503 (*n* = 11), and MCF7 alpelisib/MI-503 (*n* = 12). Tumors were harvested 22 d from treatment onset. Means ± SEM are shown. \**P* < 0.05; \*\*\*\**P* < 0.0001; unpaired *t* test. (C) Tumors derived from mice in (A). Sections are stained with the indicated antibodies, randomly selected images shown. (Scale bar, 40 μM.) *n* = 3 mice/condition. (D) Lysates prepared from tumors isolated from mice in (A) and immunoblotted with the indicated antibodies. (E) Annexin V staining in MCF7 cells transfected with MLL4/KMT2D or control siRNA and treated with DMSO, alpelisib (1 uM), and/or MI-503 (4 uM) for 120h. *n* = 3 independent experiments. Data are shown as mean ± SEM. (F) Cell viability curve in MCF7 breast cancer cells reverse transfected with siControl or siKMT2D to genetically inhibit MLL4 and treated with an 8-point range of DMSO, alpelisib and/or MI-503 for 120h. *n =* 4 independent experiments. Data are shown as mean ± SEM.

### MLL4/KMT2D genetic ablation enhances PI3K/MLL1 inhibitor-driven apoptosis and reduction in proliferation

The majority of MLL inhibitors in preclinical and clinical investigation specifically target the MLL1-containing COMPASS complex, through the loss of MLL1 association with essential complex members Menin or WDR5^18,19,25^. Because H3K4 methylation is elevated in some cancer types and is associated with a poor patient prognosis^15^, MLL 1 inhibition may function as a viable therapeutic target. However, published reports suggest that the MLL enzyme MLL4/KMT2D is an AKT substrate, and its enzymatic activity is associated with resistance to the PI3K inhibitor alpelisib in HR+ breast cancer via enhancing estrogen receptor signaling^14^. To address whether the loss of MLL4 enhances the observed synergy with dual PI3K and MLL1 inhibition, we utilized siRNA to knock down MLL4 (Fig S5B). Combined PI3K/MLL1 inhibition with MLL4 knockdown further enhanced apoptosis compared to all single- and double-agent treatment groups (Fig 4E, S5C-E) in the MCF7 cell line. Dual PI3K and MLL1 inhibition in combination with MLL4 knockdown further reduced cell viability at low inhibitor doses (Fig 4F). These data suggest that enzymatic inhibition of MLL1 and MLL4, when combined with PI3K inhibition, is a promising therapeutic modality to induce cell death and produce a durable therapeutic response in PI3K-activated cancers.

## Discussion

Of the approximately 70% of breast cancers that are HR+, 40% are characterized by aberrant PI3K pathway activity. The critical role of PI3K signaling in carcinogenesis has prompted the development clinical testing of pan- and isoform-selective PI3K inhibitors in this patient population. However, the utility of PI3K-targeted therapies is limited as inhibitors of PI3K enzymes lack efficacy as monotherapies. Thus, research to define novel mechanisms by which PI3K/AKT contributes to oncogenesis and exploiting those mechanisms to develop and test viable combination therapies for the treatment of PI3K pathway-activated cancers is critical. Here, we build upon previous studies demonstrating the PI3K effector AKT regulates cellular H3K4 methylation through the phosphorylation of the H3K4 methyltransferase MLL4/KMT2D^14^ and the H3K4 demethylase KDM5A^13^ in breast cancer. AKT-mediated KDM5A phosphorylation redistributes KDM5A from the nucleus into the cytoplasm, rendering it ineffective at H3K4me2/3 demethylation^13^. In this context, elevated promoter H3K4me3 supports the transcription and ultimately the expression of cell cycle promoting genes, encouraging proliferation and oncogenesis. In contrast, AKT- or SGK2-driven phosphorylation of the H3K4 methyltransferase MLL4 attenuates its activity^14^’^26^. PI3K inhibition enhances MLL4 activity, further driving H3K4 methylation and promoting an open chromatin conformation, primarily at enhancers typically bound by the FOXA1 pioneer factor that also contain estrogen receptor binding motifs. The open chromatin conformation supports FOXA1 binding, which then recruits ER to initiate the transcription of ER-regulated genes. These disparate mechanisms of action drive biologically distinct and separable H3K4 methylation under conditions of either PI3K pathway hyperactivation or inhibition, suggesting that H3K4 methylation is downstream of the PI3K/AKT pathway, and that combined PI3K and MLL inhibition may provide therapeutic benefit. While the data presented here focuses on the potential utility of combined PI3K/MLL inhibition in HR+, PIK3CA-mutant breast cancers, we speculate that benefit may be derived in other cancers characterized by PI3K pathway activation. PI3K signaling antagonizes the activity of ER^27^ and other nuclear hormone receptors such as androgen receptor (AR)^28^, suggesting a similar therapeutic opportunity in AR-dependent, PI3K pathway hyperactivated prostate cancers given AKT-mediated regulation of MLL4/KMT2D and KDM5A in this context. However, previous reports demonstrating that AKT dysregulates H3K4 methylation through KDM5A phosphorylation in cell lines representing all molecular subtypes of breast cancer^13^ suggest potential utility of dual PI3K/MLL inhibition in hormone receptor-independent cancers. Further studies are necessary to test this hypothesis.

Since a significant limitation of PI3K inhibitors is their cytostatic nature^22^, a major attribute in defining effective therapies to pair with PI3K inhibitors is robust apoptotic induction. Herein, we demonstrate that the combined inhibition of p110α and MLL1 synergizes to induce apoptosis of PIK3CA-mutant, HR+, breast cancer cell lines (Fig 3). Dual PI3K and MLL1 inhibition reduces the clonogenicity of PIK3CA-mutant breast cancer cell lines (Fig 1), while also decreasing their proliferation (Fig 3). This synergy is enhanced by genetic ablation of MLL4. Further, we identify a compensatory feed-back loop between MLL methyltransferases and the PI3K pathway. While additional experiments are needed to understand the mechanism(s) that underlie the observed MLL feedback loop, our results suggest that MLL inhibition may hyperactivate PI3K signaling upstream of AKT. AKT hyperactivation occurs relatively quickly and is conserved amongst MLL inhibitors that exhibit divergent scaffolding and thus function through different mechanisms – through MLL1/WDR interaction, or via MLL1/Menin binding. While the possibility of MLL-inhibitor off-target effects cannot be eliminated, AKT hyperactivation following the use of multiple MLL inhibitors suggests that compensatory AKT activation is the result of on-target MLL inhibition. The placement of MLL both upstream and downstream of AKT in the PI3K signaling network highlights the importance of dual inhibition of these enzymes as a mechanism to durably suppress PI3K pathway activation.

Our findings suggest that while PI3K and MLL1 inhibition is an effective therapeutic strategy in PIK3CA-mutant, HR+ breast cancer cell lines, this combination can be further improved. Current MLL1 inhibitors specifically target MLL1 through protein-protein interactions with essential MLL1-specific COMPASS complex members WDR5 or Menin. MLL1 methyltransferase activity is dependent on its association with WDR5; high-affinity peptidomimetic inhibitors including MM102 bind WDR5 to prevent association with MLL1^25^. Other MLL1 inhibitors including MI-136 and MI-503 share a thienopyrimidine scaffolding and function via abrogation of the MLL1/Menin association to robustly induce cell death in models of MLL1-translocated leukemias and AR-dependent prostate cancer^19,29^. While we demonstrate that these MLL1 inhibitors synergize with PI3K inhibitors in PIK3CA-mutant breast cancer, this combination does not effectively target MLL4/KMT2D, which is a downstream PI3K effector that is essential in mediating resistance to ER-degraders such as fulvestrant^14^. Because MLL4 or pan-MLL inhibitors that target the enzymatic activity of the SET-domain have only recently been published^30^ and are not commercially available, we genetically ablated MLL4 in combination with PI3K/MLL1 dual inhibition and found that knockdown of MLL4 further enhances the PI3K/MLL1-driven apoptosis (Fig 4). The results presented here suggest a therapeutic opportunity to design and test MLL4 or pan-MLL inhibitors in PI3K-activated, HR+ breast cancers.

## Materials and Methods

### Cell lines, inhibitors, siRNA reagents

MCF7, T47D, and 293T cells were maintained at 37C and 5% CO_2_ in RPMI or DMEM + 10% FBS and 1% Pen-Strep. GDC-0941, BYL719, GDC-0032, GDC-0068 were purchased from Selleck, and MI-136, MI-503, and MM-102 were purchased from Cayman. siRNAs (Silencer Select) were purchased from Invitrogen (Negative control #1, 4390843; siKMT2D, s528766).

### Cell / tumor lysis and immunoblotting

Cells were lysed in IP buffer (20 mM TrisHCl [pH 7.5], 150 mM NaCl, 5 mM MgCl2, 1% NP-40) and tumors were lysed in RIPA buffer (10 mM TrisHCl [pH 8.0], 1 mM EDTA, 0.1 mM EGTA, 1% TritonX-100, 0.1% Sodium Deoxycholate, 0.1% SDS, 140 mM NaCl), supplemented with protease and phosphatase inhibitors. Histones were acid extracted from cells by lysing in triton extraction buffer (TEB; PBS, 0.5% Triton X-100) supplemented with protease inhibitors. Tumors were sonicated (Diagenode) prior to lysis or histone extraction. Cell and tumor lysates were centrifuged 6,500 x g and histones were acid extracted from the resulting pellet with 1:1 TEB:0.8 M HCl. Histones were centrifuged and the supernatant precipitated with the addition of an equal volume of 50% TCA, then 12,000 x g centrifugation. Histones were washed one time in ice cold 0.3 M HCl in acetone and two times ice cold in 100% acetone before drying and resuspended in 20 mM Tris-HCl (pH 8.0), supplemented with protease inhibitors. Whole cell lysate or acid-extracted proteins were separated using SDS-PAGE. Proteins were transferred to nitrocellulose membranes and blocked in TBST + 5% milk. Proteins of interest were visualized and quantified after primary antibody incubation (Odyssey, Li-Cor). Primary antibodies used were as follows: AKT, AKTS473, AKTT308, S6, S6S240/44, (CST), β-actin and KMT2D/MLL4 (Millipore), and H3, H3K4me3, H3K4me2, H3K4me (Abcam).

### Cell Viability, Apoptosis and Clonogenic assays

For resazurin viability assays, 3,000 cells/well were seeded in 96-well plates. The next day, serially diluted inhibitors were added and incubated for 5 d, after which Resazurin Sigma) was added to a final concentration of 0.1 mg/ml and incubated for 5 h prior to measuring the excitation/emission (544/590) (Biotek). Cells were reverse transfected (Lipofectamine 3000) with siRNAs according to manufacturer’s instructions (Invitrogen) where indicated. To assay for apoptosis, cells were seeded and treated with the indicated inhibitors for 5 d, after which cells were detached using accutase (Biolegend) and stained using Annexin V/PI following manufacturer’s protocol (BioLegend). Cells were kept in the dark until assessment and were analyzed by flow cytometer (BDFACSymphony A3) within 1 h. To measure cell clonogenicity, 500 cells/well were seeded in a 6-well plate, and treated the following day with the designated inhibitors, and incubated for a total of 19 d with media and inhibitor change every 3-4 d, after which cells were fixed and crystal violet stained. Plates were imaged, destained, and crystal violet quantitated at OD 595 (Biotek).

### Synergy Calculation

Bliss and Loewe synergy scores were calculated in Python using the *synergy* package: https://pypi.org/project/synergy; source code https://github.com/djwooten/synergy^31^.

### Animal studies and treatment

1 x 10^7^ MCF7 cells were resuspended in 50% Matrigel (BD) and injected into the third mammary fat pads of nulliparous NCR nude mice receiving 1 uM β-Estradiol (sigma) in drinking water, and palpable tumors were measured every 3 days. For the inhibitor studies, mice were treated daily once tumors exceeded 150mm^3^. Alpelisib (MedChemExpress) was reconstituted in 0.5% methylcellulose (Sigma) and administered by oral gavage (35 mg/kg) once daily prior to tumor isolation and preparation. MI-503 (MedChemExpress) was reconstituted in 25% DMSO, 25% PEG400, and 50% PBS and administered by intraperitoneal injection (30 mg/kg) once daily prior to tumor isolation and preparation. Tumors were isolated and prepared within 3 h of treatment. All mouse experiments were conducted in accordance with protocols approved by the Institutional Animal Care and Use Committee (IACUC) of Emory University School of Medicine (EUCM).

### Immunohistochemistry

Formalin-fixed, paraffin-embedded (FFPE) prepared tumors were sectioned (Winship Cancer Tissue and Pathology Core) and mounted. Sections were deparaffinized, hydrated, and antigens retrieved with sodium citrate. Following blocking of endogenous peroxidases, sections were blocked and incubated in primary antibody overnight (4C). Primary antibodies used were as follows: AKTS473, pS6240/44 (CST), H3K4me3, H3K4me2, H3K4me1, and Ki67 (Abcam). Sections were washed and incubated in secondary antibody, after which they were incubated with ABC (Vector Labs), DAB developed (Vector Labs), and counter-stained with hematoxylin (Vector Labs). Sections were then dehydrated and mounted for imaging (Zeiss Observer A1).

### RNA isolation, sequencing, and analysis

T47D cells were isolated using the RNeasy Isolation Kit (Qiagen) according to the manufacturer’s instruction. Preparation of RNA library and transcriptome sequencing was conducted by Novogene Co., LTD (Beijing, China) and transcript abundance was quantified using salmon (Illumina DRAGEN). Differentially expressed genes were identified using DESeq2 and genes with adjusted *p*-value < 0.05 and ļlog2(FoldChange)ļ > 2 were considered differentially expressed. GSEA analysis of the pre-ranked gene list was performed using clusterProfiler and fgsea, using the msigdb oncogenic signature gene set.

### Statistical analysis

All experiments including western blot, Resazurin viability, clonogenic, and Annexin V assays, were performed in three independent experiments unless otherwise noted. Mean ± SEM are reported unless otherwise noted. Statistical significance (p < 0.05) of differences between two groups was determined by Student’s t test.

## Data availability statement

The data generated in this study are publicly available in the Gene Expression Omnibus at GSE (in progress).

## Acknowledgements and funding sources

We thank the Spangle and Hong laboratories for helpful project discussions. This study was supported by grants from the NIH (R00 CA204601 to JMS.) and the Winship Cancer Institute of Emory University (Winship Invest$ to JMS). This study was supported in part by the Cancer Tissue and Pathology shared resource of Winship Cancer Institute of Emory University and NIH/NCI (P30CA138292).

## Author Contributions

J.F., R.B.J., M.Z., A.L.H and J.M.S. designed the research. J.F., R.B.J., K.P., M.A., M.Z., R.S.L., G.C., A.L.H. and J.M.S. performed experiments and analyzed the results. J.M.S. supervised the studies. J.M.S. wrote the manuscript with input from J.F., R.B.J., K.P., and M.A.

---

**Supplementary Figure 1.**
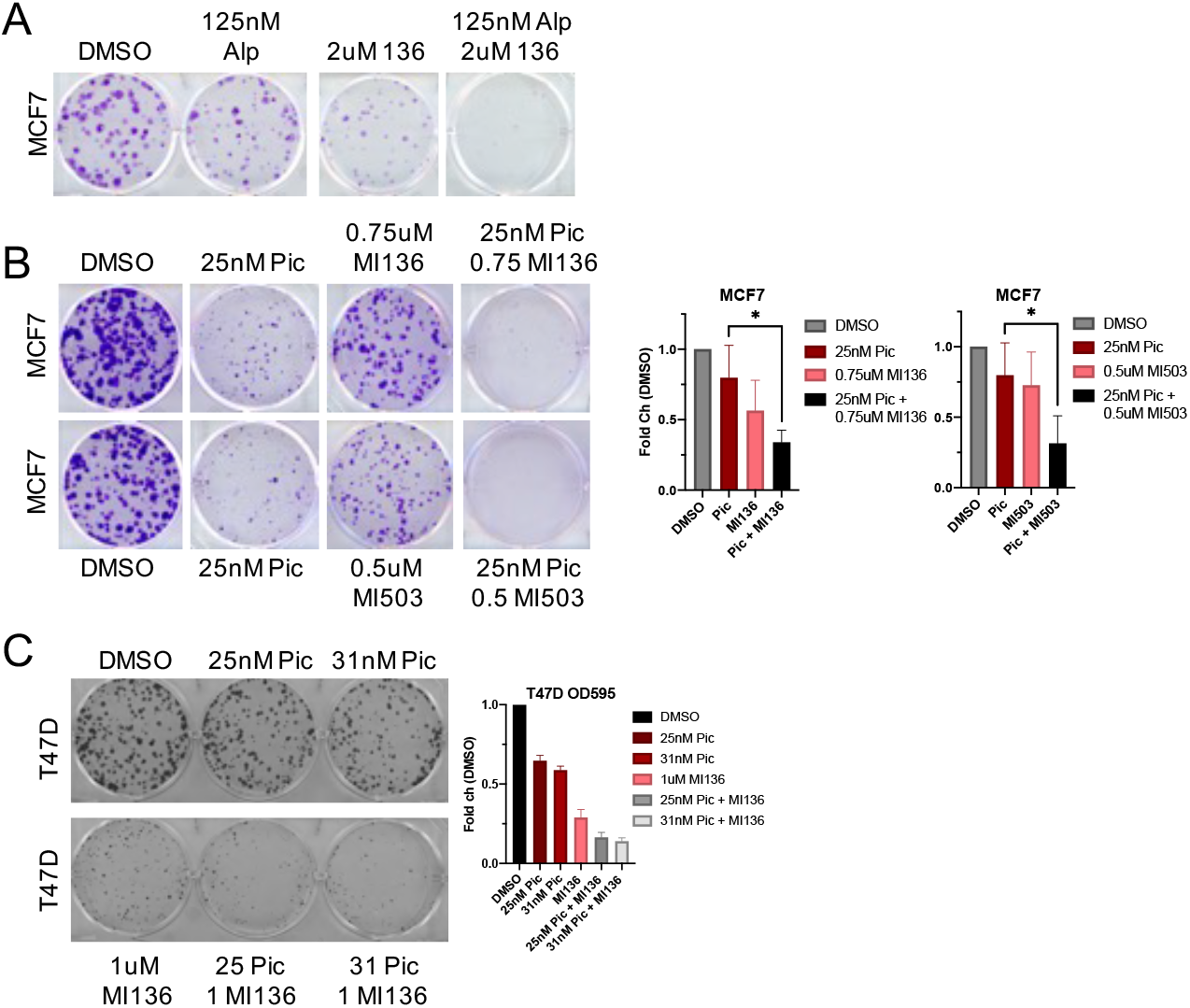
Combined PI3K and MLL1 inhibition reduces clonogenicity of HR+, PIK3CA-mutant, breast cancers. (A) MCF7 breast cancer cells treated with the indicated concentrations of alpelisib, MI-136, or DMSO for 17 days before fixation and crystal violet staining. Representative images shown. (B) MCF7 breast cancer cells treated with the indicated concentrations of pictilisib, MI-136, MI-503, or DMSO for 17 days before fixation and crystal violet staining. Representative images shown. Results shown are representative of at least 3 independent experiments. Data are shown as mean ± SEM. \**P* < 0.05, unpaired t-test. (C) T47D breast cancer cells treated with the indicated concentrations of pictilisib, MI-136, or DMSO for 19 days before fixation and crystal violet staining. Representative images shown. Results shown are representative of at least 3 independent experiments. Data are shown as mean ± SEM.

**Supplementary Figure 2.**
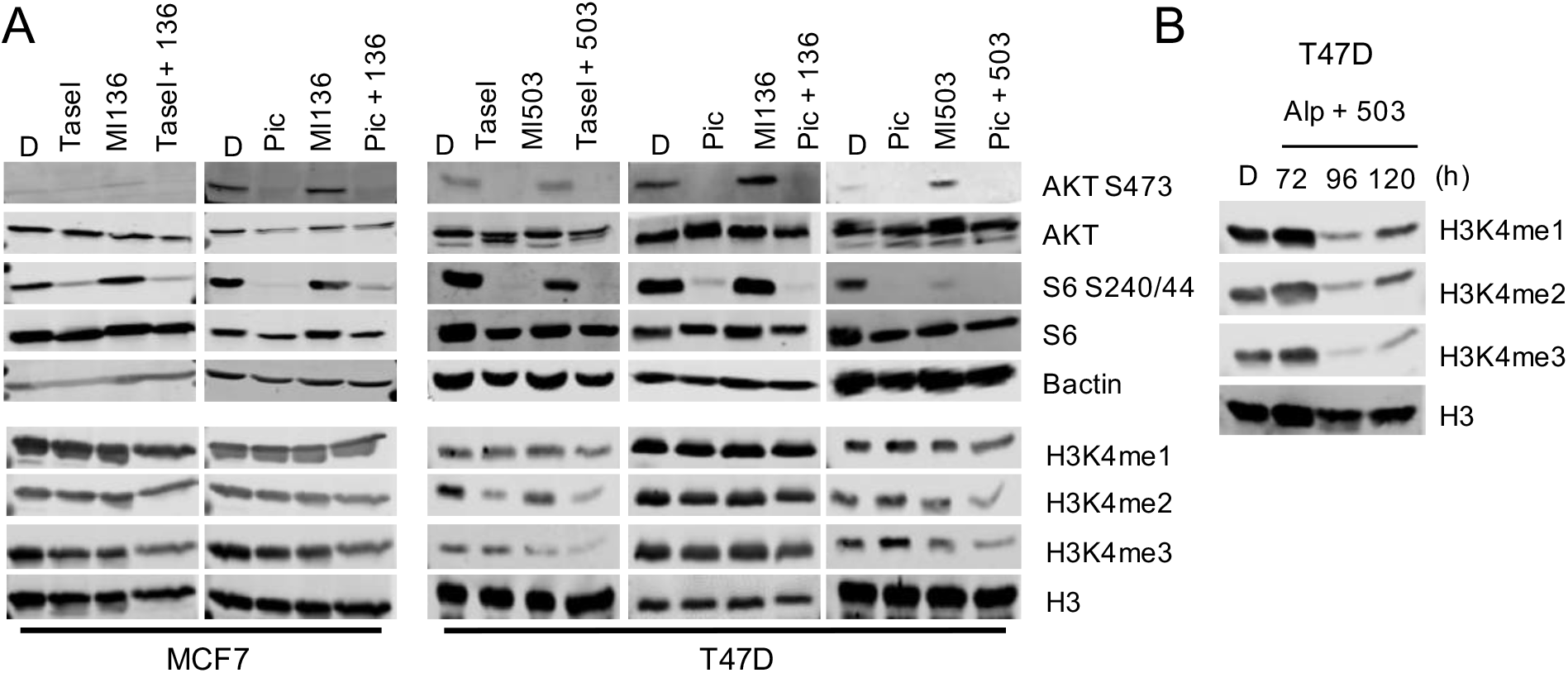
MLL1 inhibition hyperactivates AKT. (A) MCF7 (left) or T47D (right) breast cancer cells were treated with taselisib (1 uM), pictilisib (1 uM), MI-136 (4 uM), MI-503 (4 uM) or DMSO for 24h (top, PI3K effector) or 72h (bottom, histone modifications) prior to protein extraction. Lysates were immunoblotted for the indicated antibodies. (B) T47D cells treated with alpelisib (1 uM) or MI503 (4 uM) for the indicated times and lysates acid extracted and immunoblotted for the indicated antibodies.

**Supplementary Figure 3:**
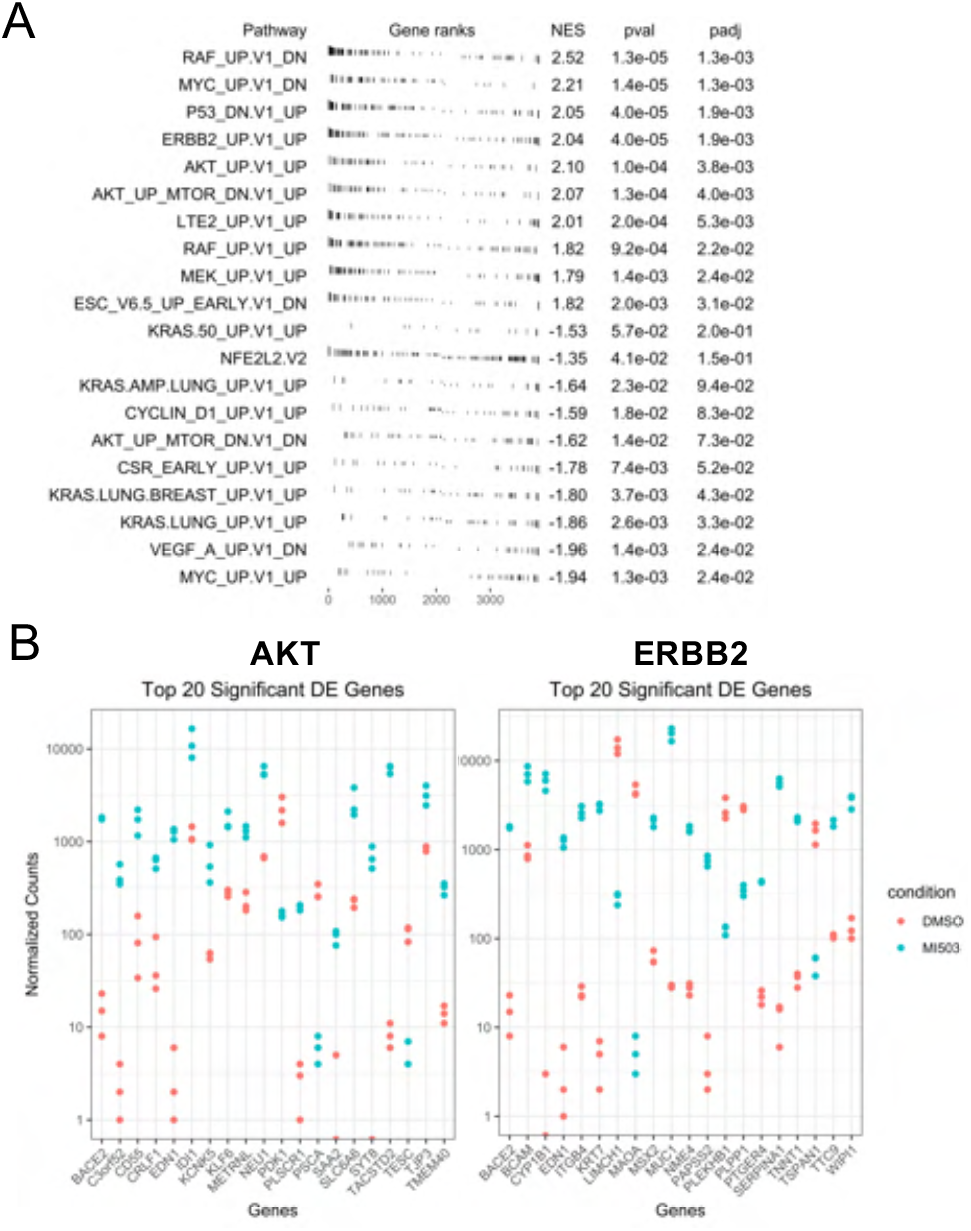
MLL1 inhibition hyperactivates PI3K effector signaling via induction of gene expression profile changes. (A). Table of enrichment graphs depicting the top 10 pathways enriched at the top and bottom of the ranked gene list, generated using the oncogenic signature gene set (c6.all.v7.4.symbols.gmt). (B) Stripplots showing differences in the normalized enrichment scores (TPM) of the top 20 genes identified as significantly enriched in the GSEA analysis between control (DMSO) and MI503.

**Supplementary Figure 4:**
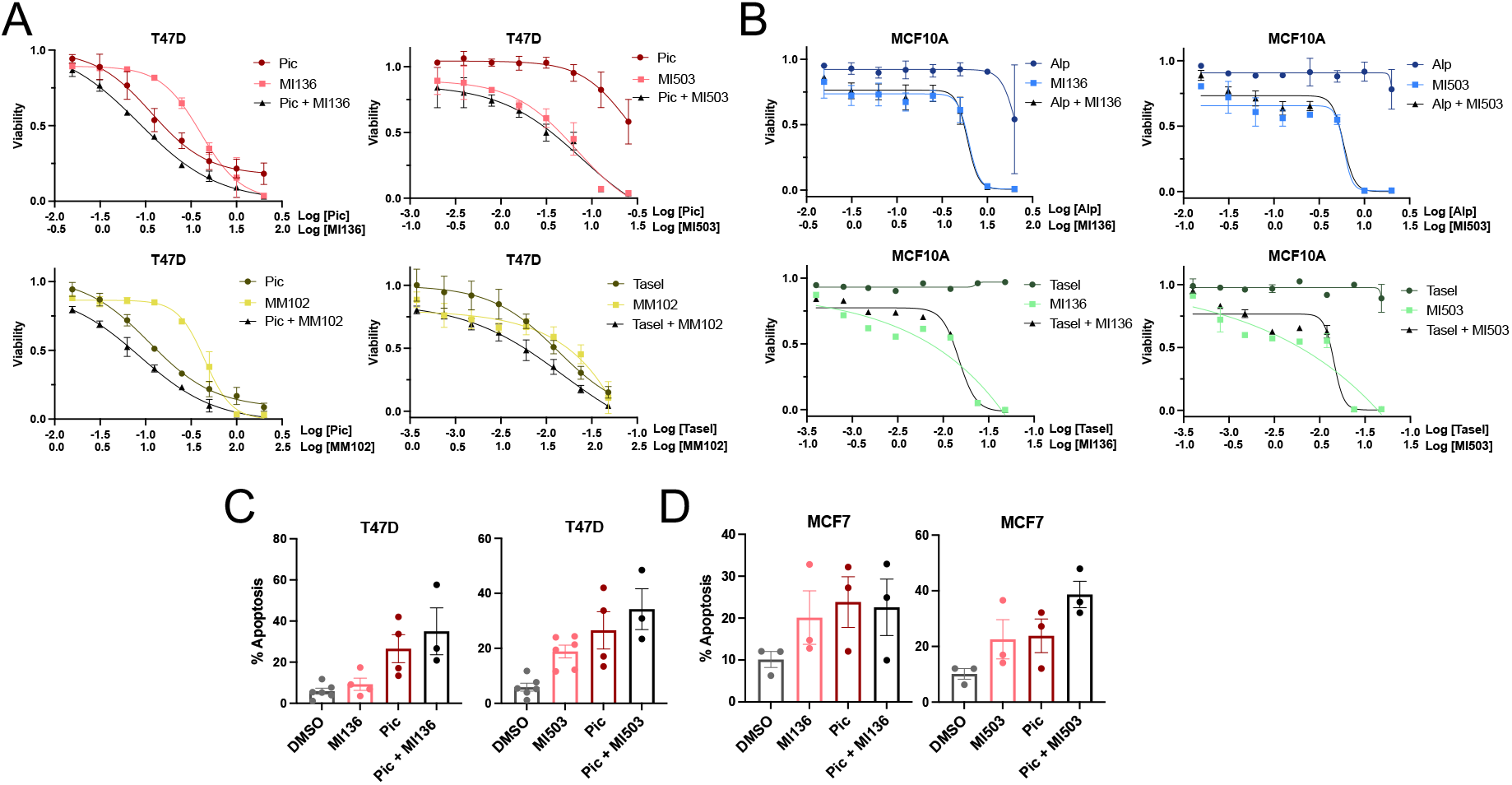
PI3K and MLL1 combination therapy synergizes to reduce viability and enhance apoptosis. (A) Cell viability curves in T47D breast cancer cells treated with an 8-point range of DMSO, pictilisib, taselisib, MI-503, MI-136 and/or MM-102 for 120h. Results shown are representative of at least 3 independent experiments. Data are shown as mean ± SEM. (B) Cell viability curves in MCF10A normal breast cells treated with an 8-point range of DMSO, alpelisib, taselisib, and/or MI-503 for 120h. Results shown are representative of at least 3 independent experiments. Data are shown as mean ± SEM. (C) Annexin V staining in T47D cells treated with DMSO, pictilisib (1 uM), MI-136 (4 uM), MI-503 (4 uM) for 120h. Data are shown as mean ± SEM; n = 3. (D) Annexin V staining in MCF7 cells treated with DMSO, pictilisib (1 uM), MI-136 (4 uM), MI-503 (4 uM) for 120h. Data are shown as mean ± SEM; n = 3.

**Supplementary Figure 5.**
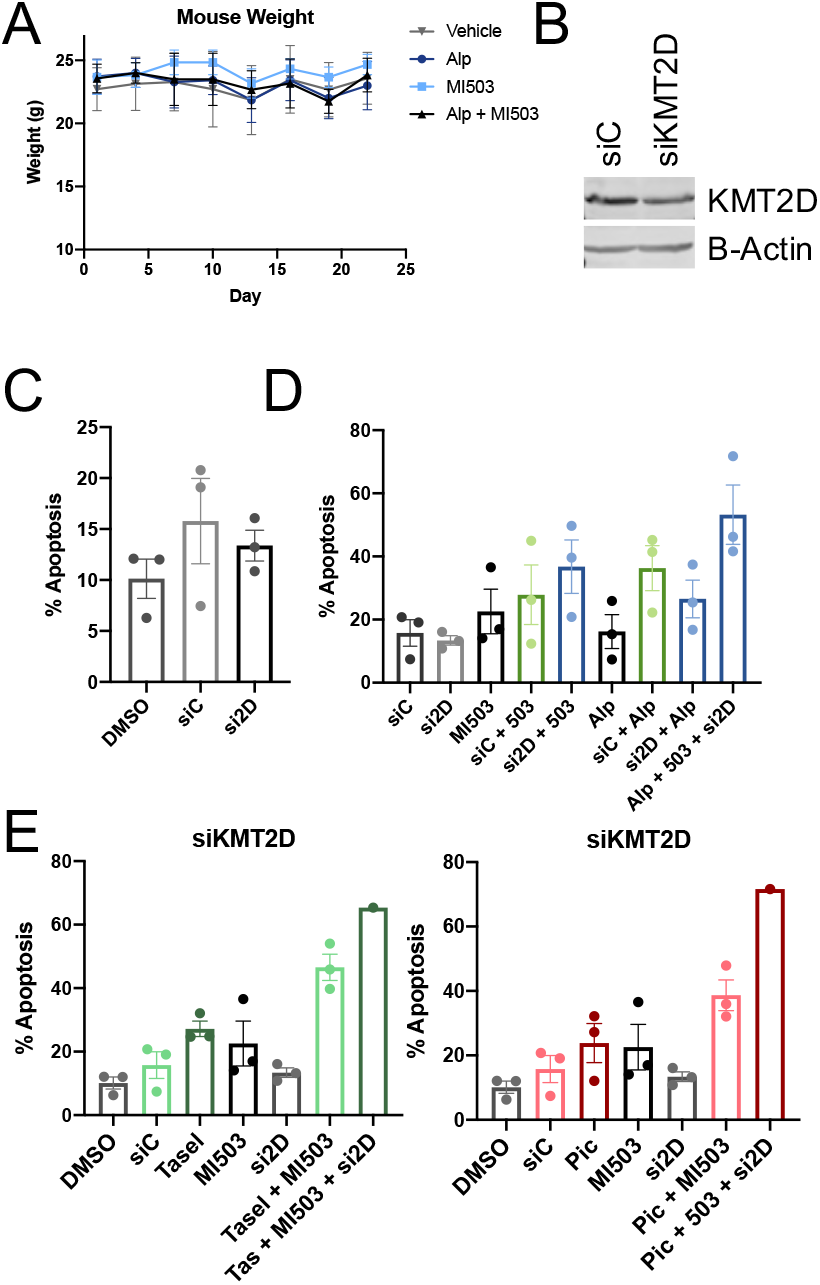
Combined MLL and PI3K inhibition provides therapeutic benefit in *in vivo* and *in vitro* models of breast cancer. (A) Mouse weight with once-daily treatment with alpelisib (45 mg/kg, gavage), MI-503 (30 mg/kg, IP), or the combination. (B) MCF7 cells reverse transfected with siControl or siKMT2D for 120h followed by lysate preparation. Lysates were immunoblotted for the indicated antibodies. (C), (D), and (E) Annexin V staining in MCF7 cells transfected with MLL4/KMT2D or control siRNA and treated with DMSO, alpelisib (1 uM), MI-503 (4 uM), taselisib (1 uM), pictilisib (1 uM) or the combination(s) for 120h. Results shown are representative of at least 3 independent experiments. Data are shown as mean ± SEM.

